# Pancreatic Duct Cells as a Potential Source for Human Islet Neogenesis: Insights from Imaging Mass Cytometry

**DOI:** 10.64898/2026.02.07.704527

**Authors:** Rui Liang, Tengli Liu, Lanqiu Zhang, Wenmiao Ma, Huixia Ren, Shusen Wang

## Abstract

The question of whether islet neogenesis occurs in adult humans has been a subject of long-standing debate. To explore the characteristics of islet endocrine cells associated with pancreatic ducts, we employed imaging mass cytometry to examine pancreatic tissues from individuals across different age groups, including those with prediabetes or type 2 diabetes (T2D). Our analysis revealed the presence of all five pancreatic islet endocrine cell types, along with two types of non-hormone-expressing endocrine cells, located within or immediately adjacent to the ducts. These cells were most abundant in infancy, with a gradual decline observed through adulthood. Notably, ductal β cells predominated in infancy, whereas ductal α cells became more prevalent in adulthood, and significantly increased in the group aged over 60 years. Obesity further increased the ductal β cells in the subjects aged over 60 years. Under prediabetic and T2D conditions, an increase in all duct-related endocrine cells was observed. These findings indicate that ductal cells may serve as a reservoir for new pancreatic endocrine cells, offering potential insights into the promotion of endogenous β cell regeneration in diabetic patients.

**Highlights:**

◯ Characterization of various islet endocrine cell types related to ducts in human pancreas.
◯ The insulin-positive cells are the dominant cells among all duct-related islet endocrine cell types during the infancy period, however, the glucagon-positive cells become the dominant cells in adulthood.
◯ T2D, Obesity, and aging are involved in the increase in the number of duct-related endocrine cells.

## Introduction

The pancreatic islets mainly contain insulin-producing β cells, glucagon-producing α cells, somatostatin-producing δ cells, pancreatic polypeptide-producing PP cells, and ghrelin-producing ε cells, which cooperate with each other to ensure glucose homeostasis. It is well-established that pancreatic duct progenitor cells contribute to endocrine cell neogenesis during embryonic development. However, the potential role of ductal cells in generating new islet cells after birth remains contentious.

The understanding of β cell neogenesis after birth has been shaped by numerous findings from various models and species over several decades ^1-4^. Although lineage tracing-based studies in rodents have continuously challenged this view by suggesting that islet regeneration solely relies on the replication of pre-existing mature β cells ^5-7^, other reports have provided substantial evidence for the occurrence of islet neogenesis in the postnatal period and throughout adult life. Both rodent and human ductal cells have been demonstrated to differentiate into all pancreatic lineages, including endocrine cells, when exposed to various growth factors ^8-10^. By superimposing pregnancy on an insulin-resistant mouse model and employing genetic lineage tracing techniques, Dirice et al. demonstrated that pancreatic duct cells contribute to β cell population through differentiation/neogenesis during physiologic (pregnancy) or pathological (T2D) insulin-resistant states ^11^.

Notably, studies addressing the issue of islet regeneration mainly depend on the lineage tracing method, which, although highly powerful for tracking specific cell lineages and their progeny, has limitations such as low labeling efficiency, promoter leakage, and the inability of tissue-specific promoters for Cre expression to fully recapitulate native expression patterns ^12,13^. The choice of promoters to label cells, the efficiency of recombination, and the duration of the tracked cells may account for some of the conflicting evidence. Additionally, notable differences exist between mouse and human islets, including cytoarchitecture, vascularization, and innervation patterns ^14^, β cell turnover magnitudes ^15^, mechanisms of adaptation to stressors like pregnancy or obesity ^16,17^, and characteristics of endocrine differentiation during islet development ^18^. Therefore, it is imperative to draw conclusions regarding islet regeneration from human samples in order to gain a comprehensive understanding.

The identification of islet neogenesis in human pancreatic tissue has predominantly relied on the observation of islet hormone-positive cells within ducts, insulin/CK19 double-positive cells, or tiny clusters of islet cells (1-3 cells) as detected by immunofluorescence ^19,20^. Recently, Domínguez-Bendala group demonstrated that sorted human P2RY1^+^/ALK3^bright+^ progenitor-like ductal cells could spontaneously mature into all pancreatic lineages, including functional β-like cells, when transplanted into immunocompromised mice ^21^, further indicated that islet cell neogenesis from ductal cells existed in adult humans. However, the dynamic changes of these cells along with age increasing from infancy to adulthood and to the elderly and in obesity and T2D conditions are still unknown.

The present study aims to analyze the development of islet cells related to duct cells during human normal growth from childhood through adulthood and into advanced age and investigate whether obesity or T2D condition affects the formation of such cells. We used imaging mass cytometry (IMC) to analyze the pancreatic tissue from non-diabetic individuals aged 1 month to over 60 years as well as obesity, prediabetes, and type 2 diabetes subjects. The IMC system combines precise laser ablation with mass cytometry to facilitate highly multiplexed protein imaging *in situ*, thereby providing proteomic information that offers deep insights into cellular states and biological functions executed by various cell types within the microenvironment. With IMC data, a comprehensive map of all islet hormone-positive (insulin, glucagon, somatostatin, PP, and ghrelin) and hormone-negative but ARX or NKX6.1-positive ductal cells was generated in each group’s pancreas.

## Materials and methods

### Human pancreas procurement and processing

To control for potential confounding effects caused by variations in pancreas positions, only sections from the tail part of the pancreas were included in this study. A total of 67 pancreatic sections were selected for analysis (Table S1). The donors were categorized into infancy (0-1 years old), childhood (2-10 years old), adolescence (12-20 years old), young adulthood (23-27 years old), adulthood (30-59 years old), and elderly (≥60 years old) non-diabetic individuals (ND) aged 0.08 to over 60 years, as well as prediabetic (PreD) and T2D organ donors. HbA1c data was retrieved from the medical records of the organ donors, and prediabetes was diagnosed when the HbA1c value fell between 5.7% and 6.4% based on the guidelines from the American Diabetes Association^22^. The ND, PreD, and T2D groups from adulthood were matched for age, BMI, and sex (Table S2). Additional donor information can be found in Table S1. Pancreas tissues were fixed in 4% paraformaldehyde, embedded in paraffin, and sectioned for subsequent IMC staining analysis. Human pancreases were obtained from organ donors with informed research consents from the Human Islet Resource Center (HIRC, China), Tianjin First Central Hospital with informed research consent. The organ donation procedure follows the regulations in China. The study protocol was approved by the Ethical Committee of Tianjin First Central Hospital (Review No.: 2016N073KY).

### Antibody conjugation and titration

All antibodies in the panel were acquired from different vendors (Table S3) and initially evaluated through immunofluorescence (IF) staining on pancreas tissue. Only antibodies that exhibited expression patterns consistent with the existing literature and demonstrated a strong signal intensity were conjugated to lanthanide metals using the MaxPar X8 Multimetal Labeling Kit (Standard BioTools), following the manufacturer’s instructions. Antibodies were diluted with Antibody Stabilizer containing 0.05% NaN3 (CANDOR® Bioscience, 131050). Serially diluted antibodies were applied to separate sections of the human pancreas to determine the optimal concentrations (Table S4).

### Tissue staining

Pancreas sections tail part were stained with our full antibody panel (Table S4). Slides were incubated for 1 hr at 70 °C in a dry oven, deparaffinized in fresh Xylol, and rehydrated through a graded alcohol series. Antigen retrieval was performed in a microwave oven for 15 min at 95 °C and naturally cool down for 20 min in Tris-EDTA, pH 9.2. After sequentially being rinsed with ddH2O and DPBS, slides were permeabilized with 0.5% triton-100 in D-PBS for 15 min, blocked with 3% BSA in D-PBS for 45 min at room temperature and then incubated with an antibody cocktail (Table S4) containing all antibodies overnight at 4 °C. The next day, after twice washing in D-PBS containing 0.2% triton-100 and twice with D-PBS, slides were counterstained with DNA stain Ir at room temperature for 45 min, washed with ddH2O, and then air dried for 20 min before the image acquisition.

### Image acquisition

Following Standard BioTools’ operation procedure and daily tuning we acquired the IMC images at a laser frequency of 200 Hz following manufacturer’s instruction. We adhered to consistent criteria for selecting regions of interest (ROIs) across all groups. Initially, H&E or immunofluorescence staining was performed on the slide of each sample, followed by IMC staining on sequential sections. ROIs were chosen based on the location of islets identified in the H&E or immunofluorescence staining results. Each ROI encompassed 800 µm×800 µm and aimed to include as many islets as possible, with at least one islet per area. On average, 9 (range 5-12) ROIs were selected per slide. Finally, we converted the MCD files to tiff images using Standard BioTools’s MCD viewer.

### Quantification of new islet cells in or attached to pancreatic ducts

Only insulin, glucagon, somatostatin, and pancreatic polypeptide and ghrelin positive cells or hormone-negative but ARX or NKX6.1 positive cells in or attached to pancreatic ducts were counted. Due to their sporadic presence, these cells were manually quantified in a blind fashion. It’s worth mentioning that only scattered endocrine cells (single cell or a minicluster with less than 3 cells), but not the big cell clusters, in duct were considered in order to exclude the possibility that small clusters are mature islets adjacent to duct.

### Statistics

Figure drawing and data processing were conducted using GraphPad Prism v9.0 (GraphPad Software, La Jolla, CA, USA). Student’s t test was used for analyzing the group differences. p<0.05 were considered statistically significant.

## RESULTS

### Characterization of duct-related islet endocrine cells in human pancreases

We examined the pancreas sections from various age groups to identify islet endocrine cells associated with duct cells, referred to as duct-related endocrine cells (Figure 1). These sections showed the presence of all islets endocrine cell types, including insulin, glucagon, somatostatin, PP, and ghrelin-positive cells, either within or closely adjacent to the ducts (Figure 2A & 2B). ARX and NKX6.1 are key transcription factors associated with the specification and differentiation of α and β cells, respectively. Here, hormone-negative but ARX or NKX6.1-positive cells were also found within the duct lines across all age groups (Figure 2A&2B). According to the differentiation trajectory during the embryonic stage, these cells may represent early-stage endocrine cells that have committed to a specific endocrine cell fate but have not yet begun hormone expression. Interestingly, 62.6% of the hormone-negative ductal endocrine cells expressed CK19, while only 2.3% of the hormone-positive ductal endocrine cells were CK19-positive (Figure S1). The difference in CK19 expression ratios between these two groups suggests a potential conversion process, wherein CK19-expressing ductal cells may transform into CK19-negative islet endocrine cells.

**Figure 1.**
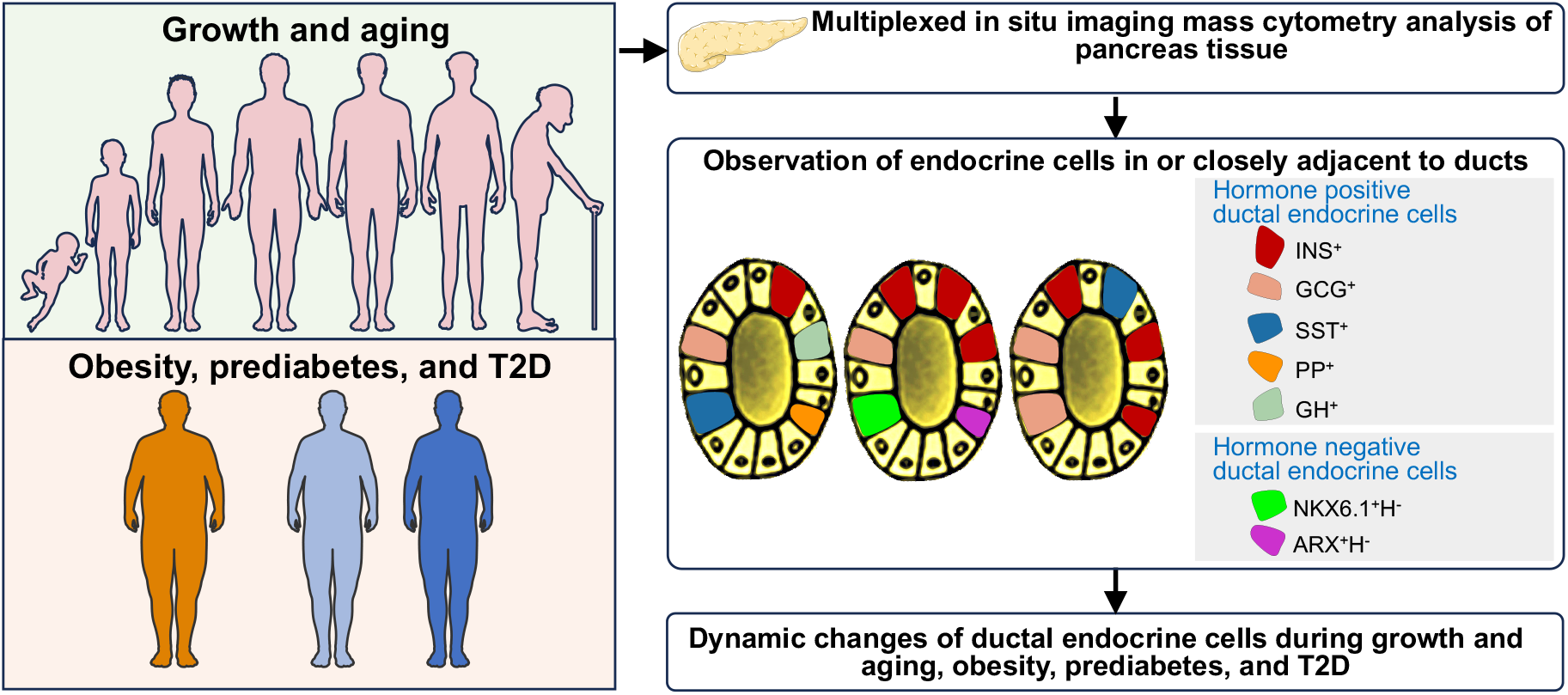
Schematic of study overview. INS^+^: insulin expressing ductal cells, GCG^+^: glucagon expressing ductal cells, SST^+^: somatostatin expressing ductal cells, PP^+^: pancreatic polypeptide expressing ductal cells, GH^+^: ghrelin expressing ductal cells, NKX6.1^+^H^-^: NKX6.1 positive but pancreatic hormone negative ductal cells; ARX^+^H^-^: ARX positive but pancreatic hormone negative ductal cells, T2D: type 2 diabetes.

**Figure 2.**
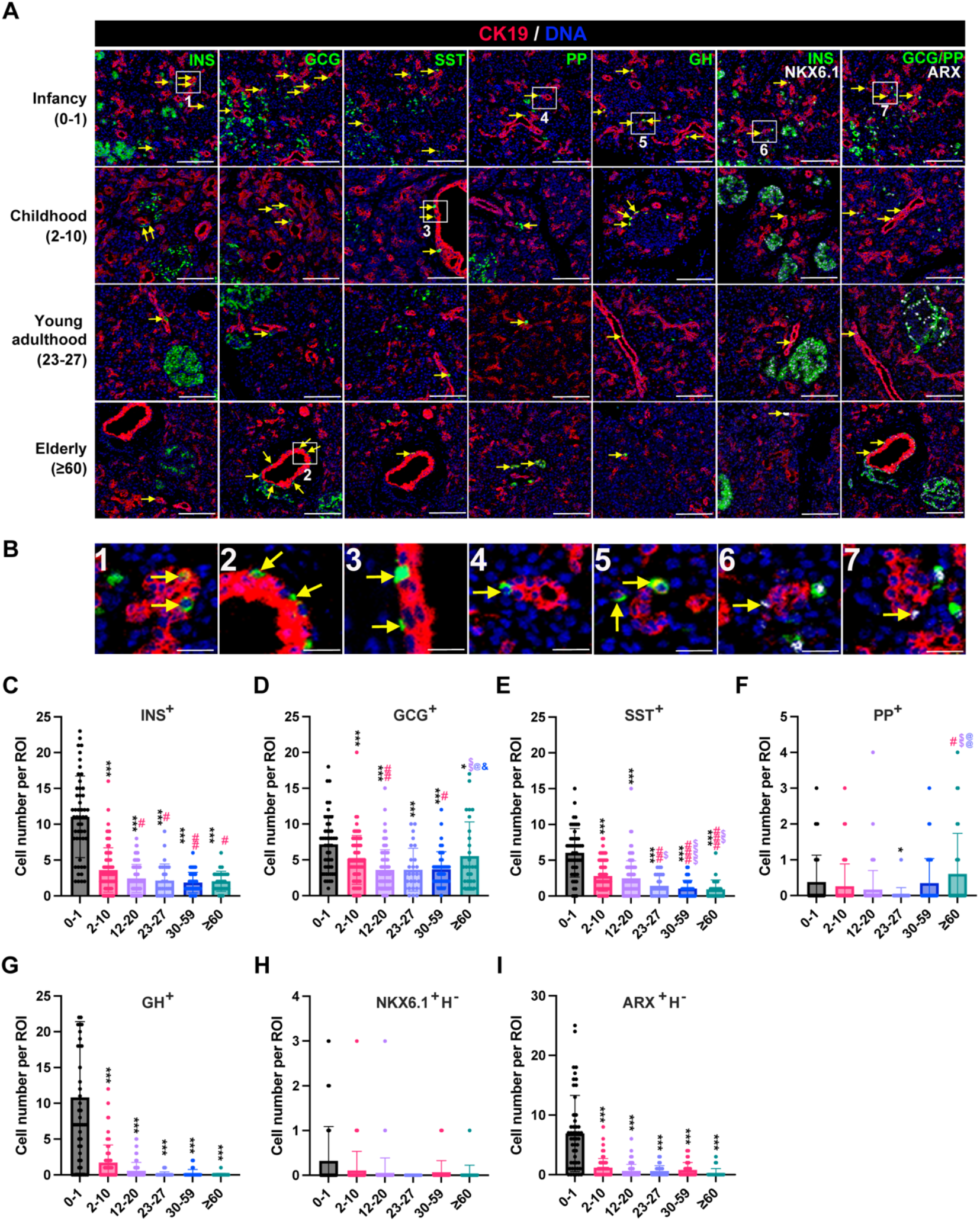
Characteristics of duct-related endocrine cells in human pancreases sections from different age groups. (A) Representative images of duct-related endocrine cells, including hormone-positive cells (insulin, glucagon, somatostatin, PP, and ghrelin-positive cells) and hormone-negative but ARX or NKX6.1-positive cells. Scale bar: 100 µm. The yellow arrow represents duct-related endocrine cells in each image. (B) Zoomed images of representative cell types. Scale bar: 20 µm. The yellow arrow represents duct-related endocrine cells in each image. (C-I) Quantification of cell number of each duct-related endocrine cell type per ROI in different age groups, including INS^+^ (C), GCG^+^ (D), SST^+^ (E), PP^+^ (F), GH^+^ (G), NKX6.1^+^H^-^ (H), and ARX^+^H^-^ (I). * p < 0.05, ** p < 0.01, *** p < 0.001 vs 0-1 years old group; # p < 0.05, ## p < 0.01, ## p < 0.001 vs 2-10 years old group; $ p < 0.05, $$ p < 0.01, $$$ p < 0.001 vs 12-20 years old group; @ p < 0.05, @@ p < 0.01, @@@ p < 0.001 vs 23-27 years old group; & p < 0.05 vs 30-59 years old group.

### The highest number of ductal-endocrine cells was observed in the pancreas during infancy

In infancy, the highest number of these duct-related hormone-positive cells, except for PP-positive cells, was observed (Figure 2A-2B). The quantity of these cells significantly decreased after infancy, continued to decline throughout childhood, and then stabilized during adolescence (Figure 2C-2G). Among the ductal endocrine cells in infancy, β cells were the most prevalent, followed by ε cells and α cells. Ductal PP cells were consistently rare across all age groups (Figure 2F).

The hormone-negative NKX6.1-positive cells were relatively scarce in all age groups (Figure 2H). Conversely, the highest number of hormone-negative ARX-positive cells was observed in infancy, with a steep decline thereafter (Figure 2I).

In individuals over 60 years of age, the number of glucagon-positive cells and PP-positive cells significantly increased compared to those in other adulthood and adolescent groups (Figure 2D), while no significant changes were observed in other cell types in the elderly.

### All duct-related endocrine cells increased in prediabetes and T2D subjects

We further investigated the effect of prediabetes and T2D on the abundance of duct-related endocrine cells in the human pancreas (Figure 3A). Prediabetes was defined by their HbA1c between 5.7-6.4%^22^. The results demonstrated that the number of duct-related insulin, glucagon, somatostatin, PP-positive cells, as well as hormone-negative ARX or NKX6.1-positive cells, were significantly increased under both prediabetic and T2D conditions (Figure 3B-H). This suggests that specific islet hormone promoters may be activated in certain ductal cells, leading to the generation of new islet cells. These dynamic changes challenge the idea that these cells in the adult pancreas are mere remnants of embryonic endocrine development, suggesting that the ductal endocrine cell formation can be stimulated by prediabetes and T2D conditions.

**Figure 3.**
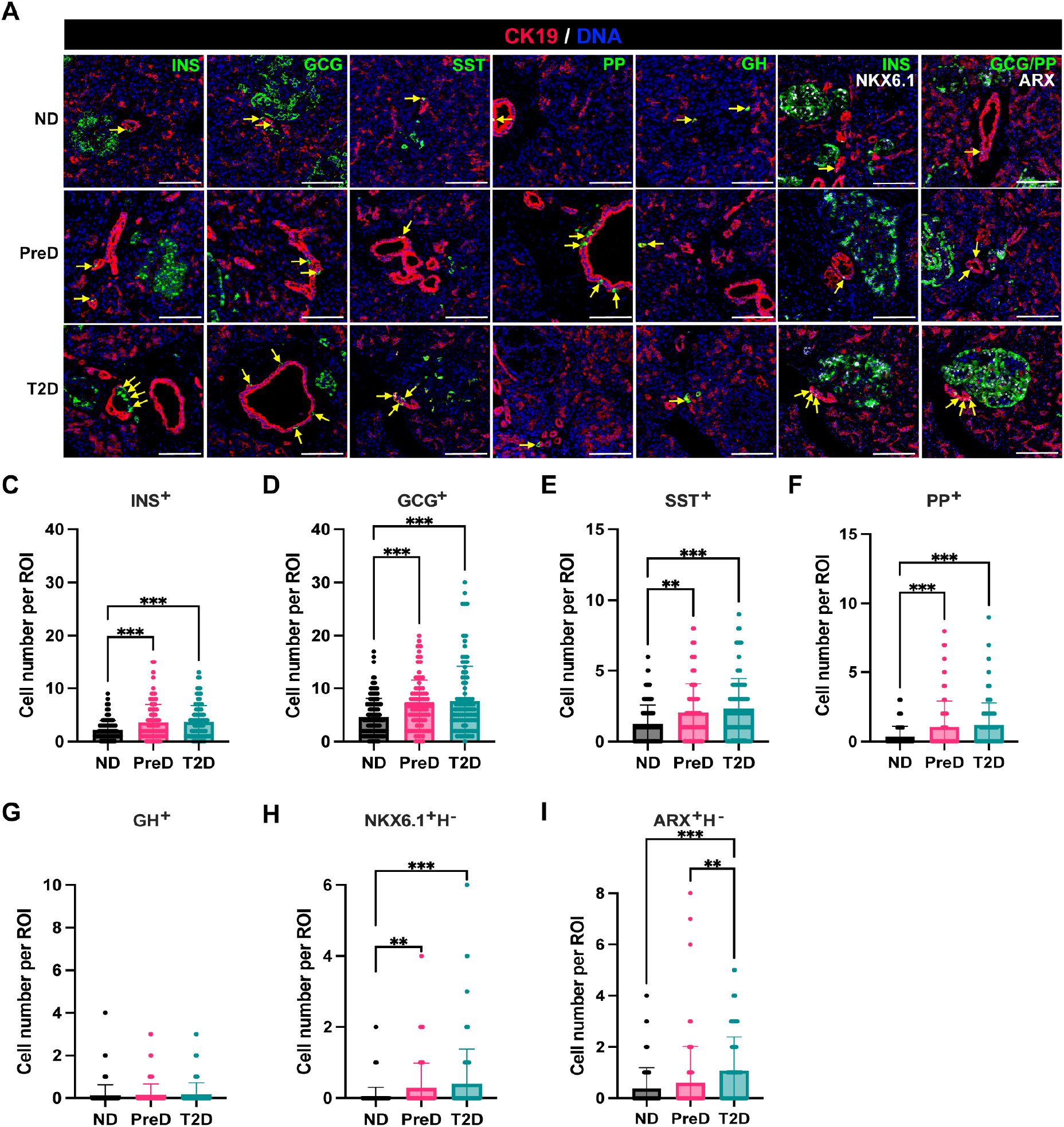
Number of duct-related endocrine cells in ND, PreD, and T2D subjects. (A) Representative images of duct-related endocrine cells, including hormone-positive cells (insulin, glucagon, somatostatin, PP, and ghrelin-positive cells) and hormone-negative but ARX or NKX6.1-positive cells in ND, PreD, and T2D. Scale bar: 100 µm. The yellow arrow represents duct-related endocrine cells in each image. (B-H) Quantification of cell number of each duct-related endocrine cell type per ROI in ND, PreD, and T2D, including INS^+^ (B), GCG^+^ (C), SST^+^ (D), PP^+^ (E), GH^+^ (F), NKX6.1^+^H^-^ (G), and ARX^+^H^-^ (H). ** p < 0.01, *** p < 0.001.

### Increase in duct-related β cells in obese aging subjects

In this study, adults aged 30 to 59 years and older adults over 60 years were stratified into obese (BMI >25) and lean (BMI ≤ 25) subgroups. Subsequent analysis revealed a higher abundance of duct-related insulin-positive cells in the obese subgroup of the elderly, while no significant differences were observed in any endocrine cell type between the obese and lean subgroups within the 30-59-year-old group (Figure 4A-G). These results suggest that obesity has a greater impact on ductal endocrine cell formation in the elderly than in younger subjects.

**Figure 4.**
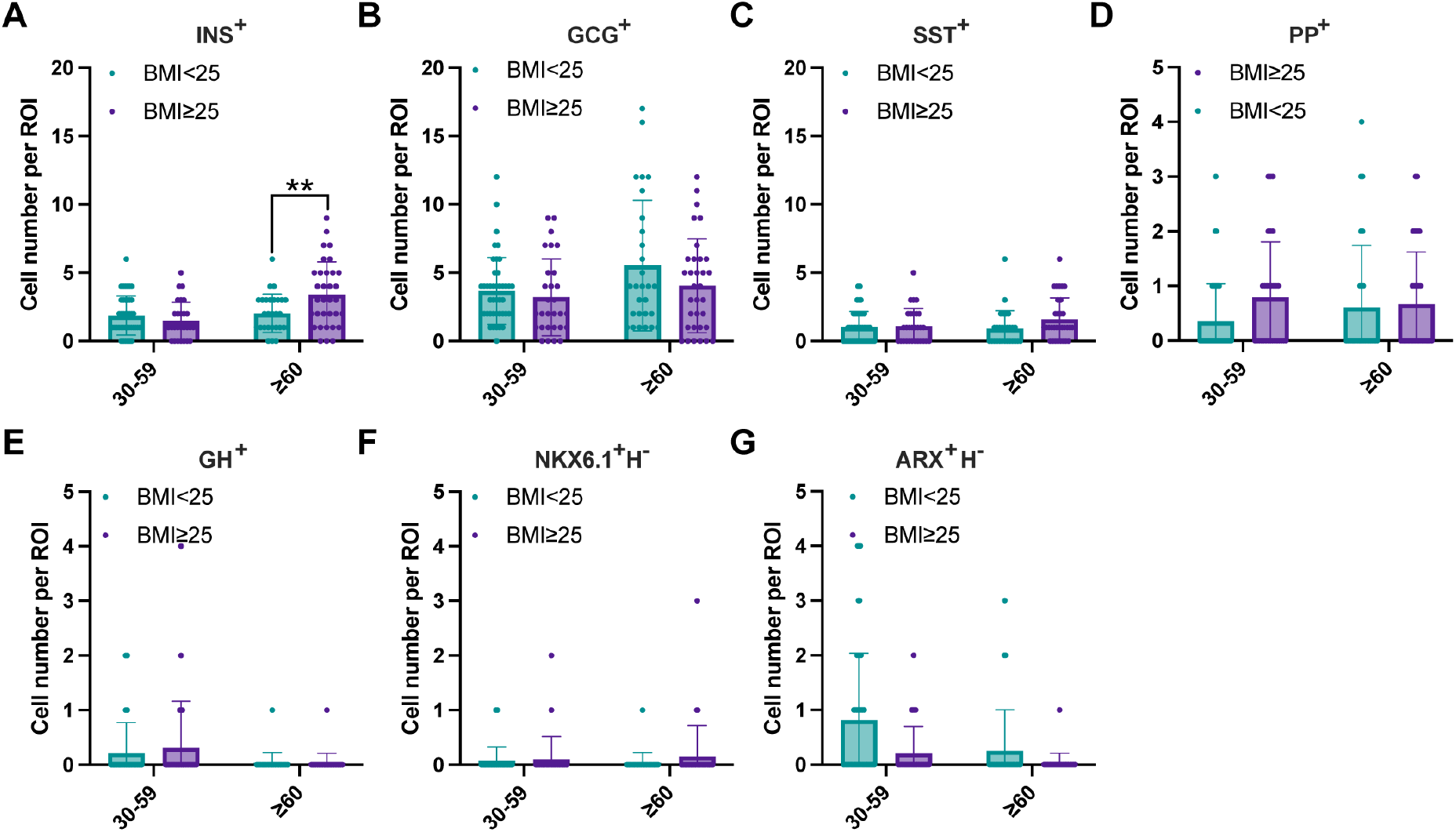
The number of duct-related endocrine cells from the subjects with or without obesity. (A-G) Quantification of cell number of each duct-related endocrine cell type per ROI in subjects with or without obesity, including INS^+^ (A), GCG^+^ (B), SST^+^ (C), PP^+^ (D), GH^+^ (E), NKX6.1^+^H^-^ (F), and ARX^+^H^-^ (G). ** p < 0.01

### α cells as the predominant duct-related endocrine cells in adults

Another striking observation in our study is the shift in the dominant duct-related endocrine cell types. During infancy, ductal β cells represented the highest proportion of endocrine cells among all ductal-related endocrine cells, accounting for 33% of all duct-related endocrine cells (Figure 5A). Ductal α cells, following the ghrelin-positive cells, were the third most abundant cells (Figure 5A). However, after infancy, the fraction of ductal α cells surpassed that of ductal β cells, making them the most prevalent endocrine cell type (Figure 5A). Ductal δ and PP cells were relatively rare compared to ductal β and α cells (Figure 5A). During diabetes development, the ductal α cell fraction was continually higher than that of ductal β cells in non-diabetes, prediabetes, and T2D subjects (Figure 5B). Our results suggest that pancreatic ducts are more prone to generate α cells in human adulthood.

**Figure 5.**
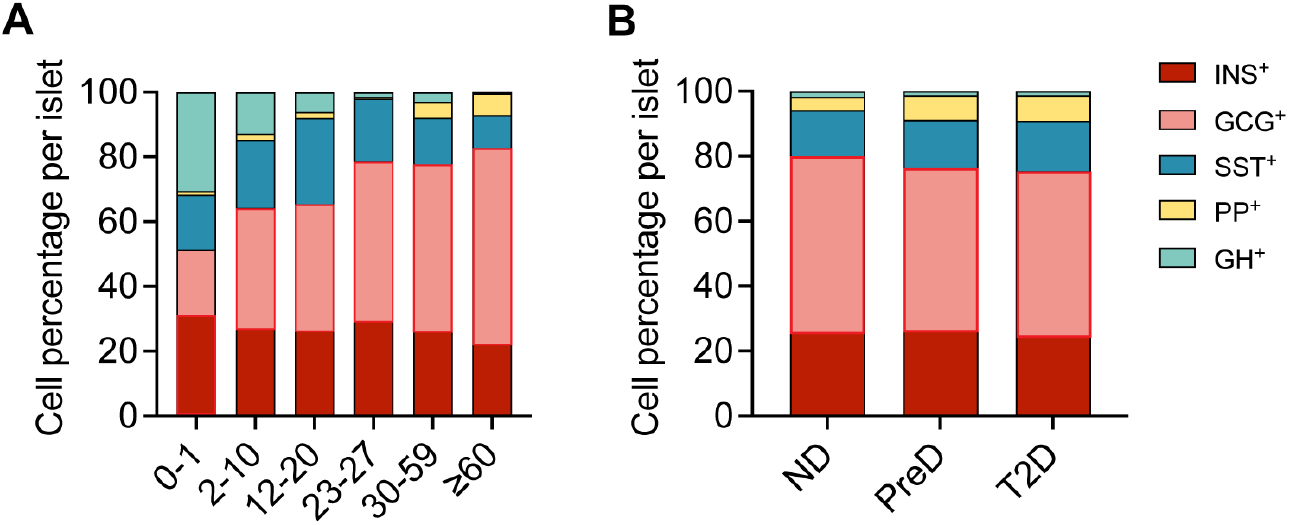
Changes in the dominant duct-related endocrine cell types. (A) Fractions of five hormone-expressing duct-related endocrine cell types in human pancreas sections from different age groups. (B) Fractions of five hormone-expressing duct-related endocrine cell types in ND, PreD, and T2D human pancreas sections. Red outlines indicate the most abundant cell type in each column.

## DISCUSSION

In this study, we utilized IMC to conduct a comprehensive analysis of duct-related endocrine cells across various age groups and disease conditions. Leveraging the advantages of highly multiplexed tissue imaging, we successfully captured all islet hormone-positive and hormone-negative ARX or NKX6.1-positive cells associated with ducts in each age group.

### Ductal cells are a source of β cell neogenesis after birth and continually contribute to replenishing endocrine cells in adult humans

Whether β cell neogenesis contributes to postnatal β cell mass increase and their origins remains unclear. In our study, we used CK19 as a marker for ductal cells, and CK19 antibody staining results revealed classic pancreatic duct cell morphology, consistent with previous IMC studies in humans^23^. We marked pancreatic endocrine cells using various hormone expressions and key transcription factors. Notably, we discovered a significant presence of pancreatic endocrine cells either within or closely adjacent to ducts. Importantly, many of these cells, particularly those still hormone-negative, co-expressed the ductal cell marker CK19. This finding contrasts with a study in mice, where pancreatic exocrine duct cells were reported to give rise to insulin-producing β cells during embryogenesis but not after birth, likely due to species differences^24^. Additionally, we observed that during infancy, some islets are located very close to the ducts, or even surrounded by them (Figure S2). Small islet cell clusters adjacent to or on the duct lining were also observed across different groups (Figure S3). These results together indicated that ductal cells contribute to postnatal endocrine cell formation.

The ongoing contribution of ductal cells to the replenishment of endocrine cells in adult humans remains poorly understood. In our study, we observed a steep decline in ductal-related endocrine cells after infancy, with levels remaining low in adults. Under non-diabetic conditions, duct-related endocrine cells remained relatively stable, with the exception of an increase in ductal α cells in lean subjects over 60 years old and an increase in ductal β cells in obese subjects over 60. Under pathological conditions such as prediabetes and type 2 diabetes (T2D), all endocrine cell types associated with ducts showed significant increases. These physiological and pathophysiological conditions share common factors like inflammation, peripheral insulin resistance, accumulation of senescent cells, and metabolic stress, which can lead to β cell dysfunction and insufficient insulin secretion^25-27^. Prolonged inadequate insulin secretion may stimulate compensatory β cell regeneration, potentially through neogenesis from ductal cells. Our findings align with earlier studies, such as Dirice et al., who reported an increase in insulin-positive ductal cells in T2D patients^11^. Similarly, A. Pugliese group once reported significantly elevated insulin expression in the ducts of pancreas transplant patients experiencing a recurrence of type 1 diabetes (T1D)^28^. The number of insulin-positive ductal cells was quite substantial in their study compared to ours, possibly due to the different disease backgrounds: their subjects were T1D patients with autoimmunity recurrence post-pancreas transplantation, whereas our subjects were with type 2 diabetes (T2D), or non-diabetic ones. Overall, these dynamic changes in ductal endocrine cells under different conditions suggest that these cells are not merely remnants of embryonic pancreas development.

The molecular mechanism for ductal cell to β cell transition is unknown. Previous studies have identified PDX1 and NGN3 as two of the key transcription factors for endocrine cell differentiation. In this study, we observed that PDX1 is expressed both in islet β cells and ductal cells (Figure S4), though with a higher expression level in the former, which aligns with the role of PDX1 in pancreas organogenesis, where PDX1-expressing progenitors give rise to both exocrine and endocrine lineages. NGN3-positive ductal cells were also observed in some cases (Figure S5), suggesting that NGN3 may play a role in regulating the transition from ductal cells to endocrine cells.

### α cells are the most abundant ductal-related endocrine cells in adults

What type of endocrine cells is easier to convert from ductal cells is worth investigating. In adults, a lateral comparison of each cell type fraction indicated that α cells have the highest abundance in all duct-related endocrine cell types, and the ductal α cell fraction presented an increasing trend along with increasing age. In addition, the ARX^+^GCG^-^PP^-^ duct-related endocrine cells are also more prevalent than the NKX6.1^+^INS^-^ cells. In broad agreement with these findings, Webb et al. demonstrated that there were substantial α cells emanating from the pancreatic ductal structure in a patient suffering from diabetes and severe chronic pancreatitis^29^. Furthermore, although we also observed Ki67-expressing cells in or around ducts, few overlapped with the duct-related endocrine cells, which ruled out the possibility that the increase of duct-related endocrine cells is due to the different proliferation potential of preexisting ductal endocrine cells. Although in infancy, ductal β cells are the most abundant, ductal α cells are the most prevalent in adults. These results may imply that either in the physiological condition or pathophysiological condition of human adults, there is a predisposition for α cells to regenerate from ductal cells more readily than β cells.

In addition, we observed that duct-associated α cells were increased in PreD and T2D. In T2D, a known clinical observation related to α cell functional change is fasting hyperglucagonemia ^30,31^. Whether the increase of ductal α cells contribute to the existent mature α cell pool and hence lead to elevated glucagon secretion worth further investigation.

## Limitations

Although we observed different-staged ductal endocrine cells, such as both CK19 positive and negative cells and hormone positive and negative cells, they are only snapshots of the pancreas, which is a key limitation in human pancreas study. Real-time observation of cell differentiation or transition in human pancreas tissue faces two main challenges: long-term observation and cell identity labeling. Pancreatic slice culture can facilitate long-term observation (over ten days) in an intact environment ^32-34^. For cell type identification, adenoviruses with reporter genes controlled by cell-specific promoters offer a promising solution, enabling precise tracking of native, transitioning, and transitioned cells. Combining this labeling strategy with slice culture could potentially allow real-time observation of these processes. However, the efficient combination of these technologies is still under development.

In addition, the study cohorts used in this study mainly include male subjects, with only a few female samples involved, so we were not able to stratify the samples by sex. Although acinar cells have been reported to transform to beta cells in rodents ^35-37^, this study did not incorporate antibodies specific to pancreatic acinar cell markers, so we are unable to assess whether any pancreatic acinar cells were expressing islet hormones. We did not investigate the triggering factors and signaling pathways involved in the conversion of ductal cells to endocrine cells in this study, but this is an area worthy of future research for developing strategies to stimulate endogenous regeneration. In the future, spatial transcriptomics technologies could be utilized in human pancreas studies to shed light on the cell differentiation trajectories and elucidate the underlying molecular mechanisms and signaling pathways during the ductal cell to endocrine cell transition.

## Conclusions

The increased number of hormone-positive and hormone-negative endocrine cells associated with ducts in obesity, prediabetic, and T2D subjects suggested these cells might be new islet cells generated from ductal cells. Furthermore, the characterization of these cells in each age group indicated that β cells are the dominant cell type among all new endocrine cells in infancy; however, α cells are the most prominent type in adulthood either in physiologic or T2D conditions. These findings provide evidence for islet cell neogenesis from ducts in human pancreases during different periods and disease conditions in human life.

## Acknowledgement

This work was supported by National Key Research and Development Program of China (2024YFA1109002 to S.W.), National Natural Science Foundation of China (82570940 to R.L., 82070805 to S.W.), National Key Research and Development Program of China (2020YFA0803700 to S.W.), Tianjin Municipal Health Commission (TJWJ2025QN041 to T. L.), Tianjin Key Medical Discipline Construction Project (Grant No. TJYXZDXK-3-013B to L.Z and TJYXZDXK-3-011B to S.W.), Chinese Institutes for Medical Research-Beijing to H.R., and Tianjin Municipal Human Resources and Social Security Bureau (XB202011 to Human Islet Resource Center, Tianjin). The guarantor (S.W) takes full responsibility for the work as a whole, including the study design, access to data, and the decision to publish.

## Supplementary Figures

**Figure S1.**
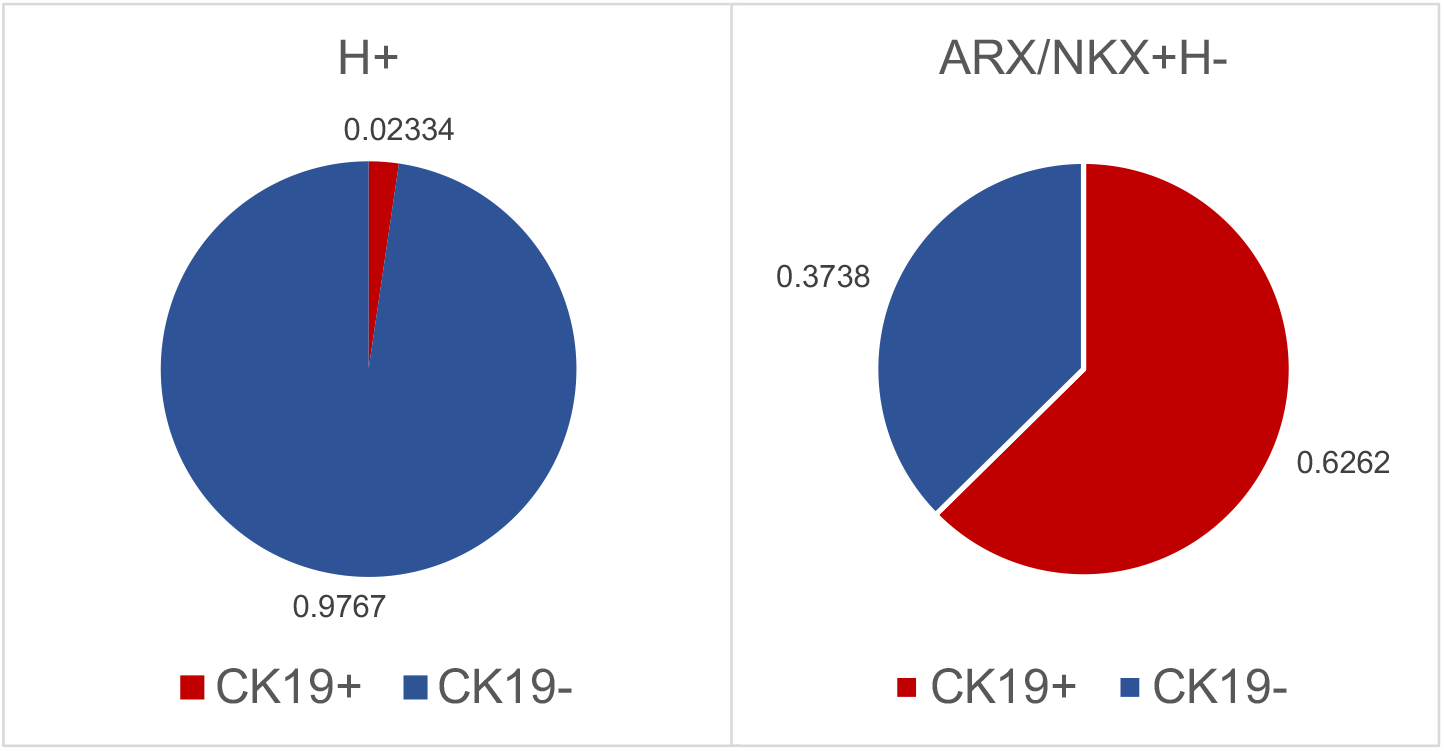
CK19 positive cell fraction in hormone-positive ductal cells and hormone-negative ductal cells.

**Figure S2.**
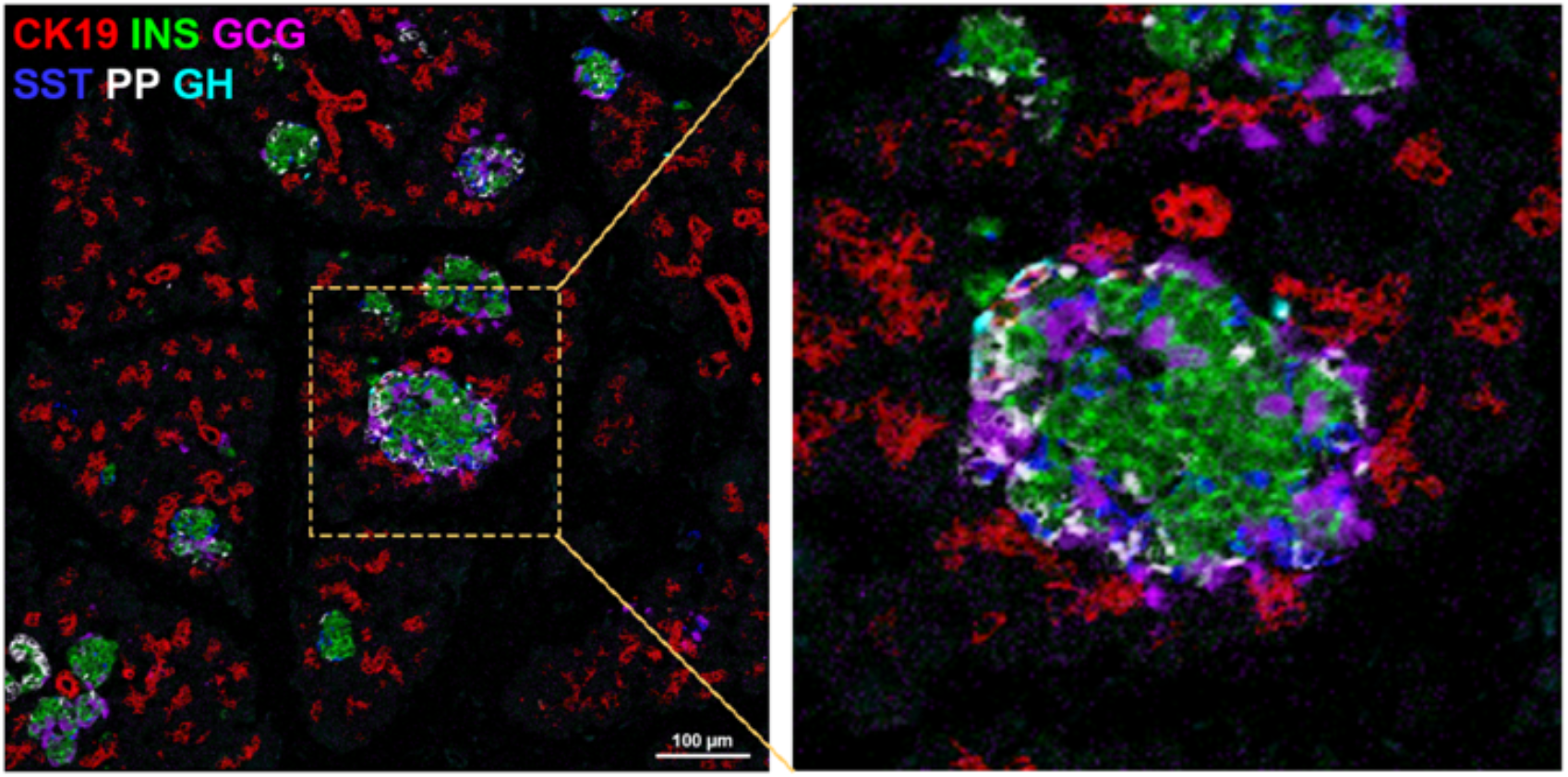
Islet surrounded by small ducts. Red, CK19; Green, INS; Purple, GCG; Blue, SST; White, PP; Cyan, GH.

**Figure S3.**
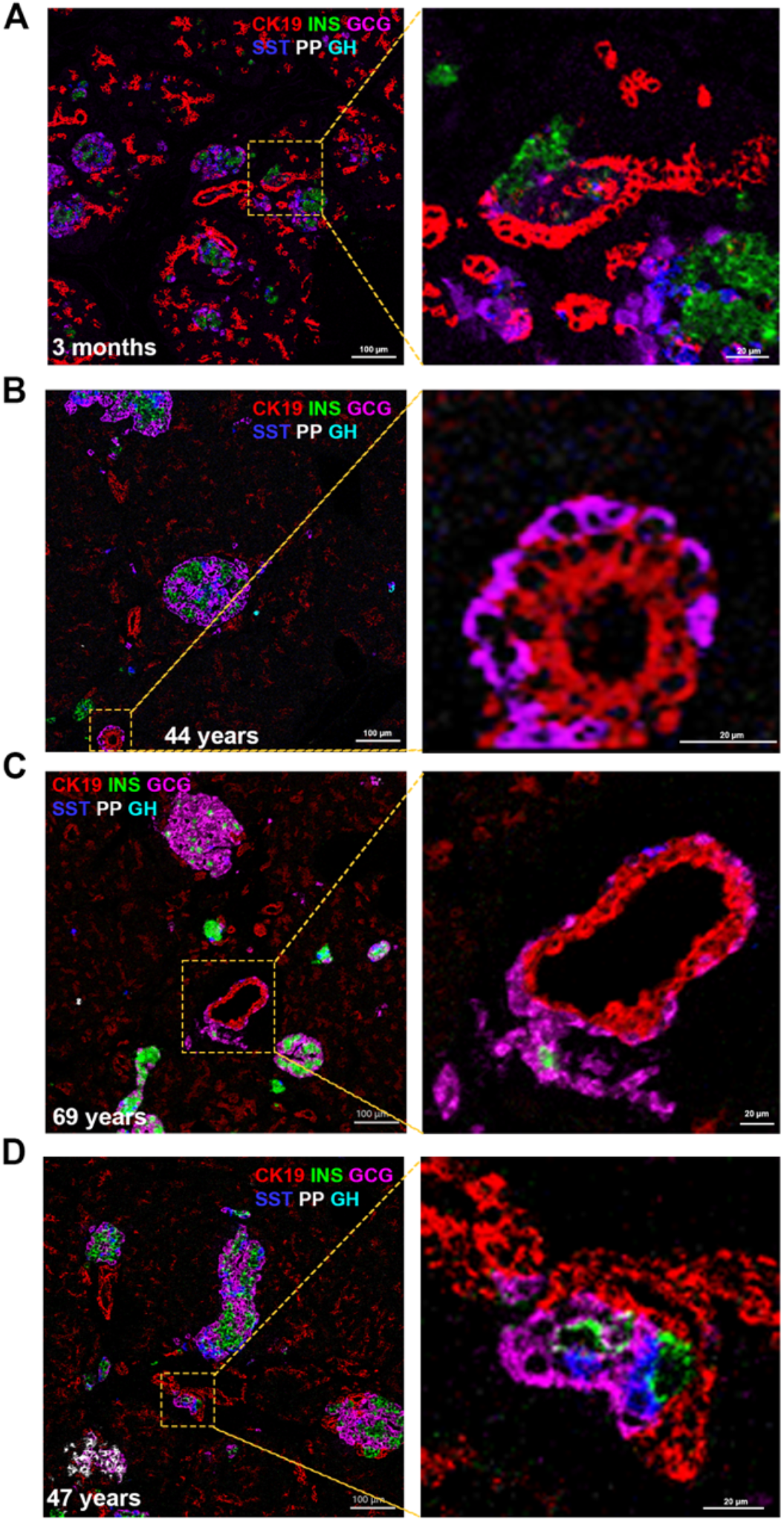
Representative IMC images showing sprouting islet cell clusters on or associated with ducts. Red, CK19; Green, INS; Purple, GCG; Blue, SST; White, PP; Cyan, GH.

**Figure S4.**
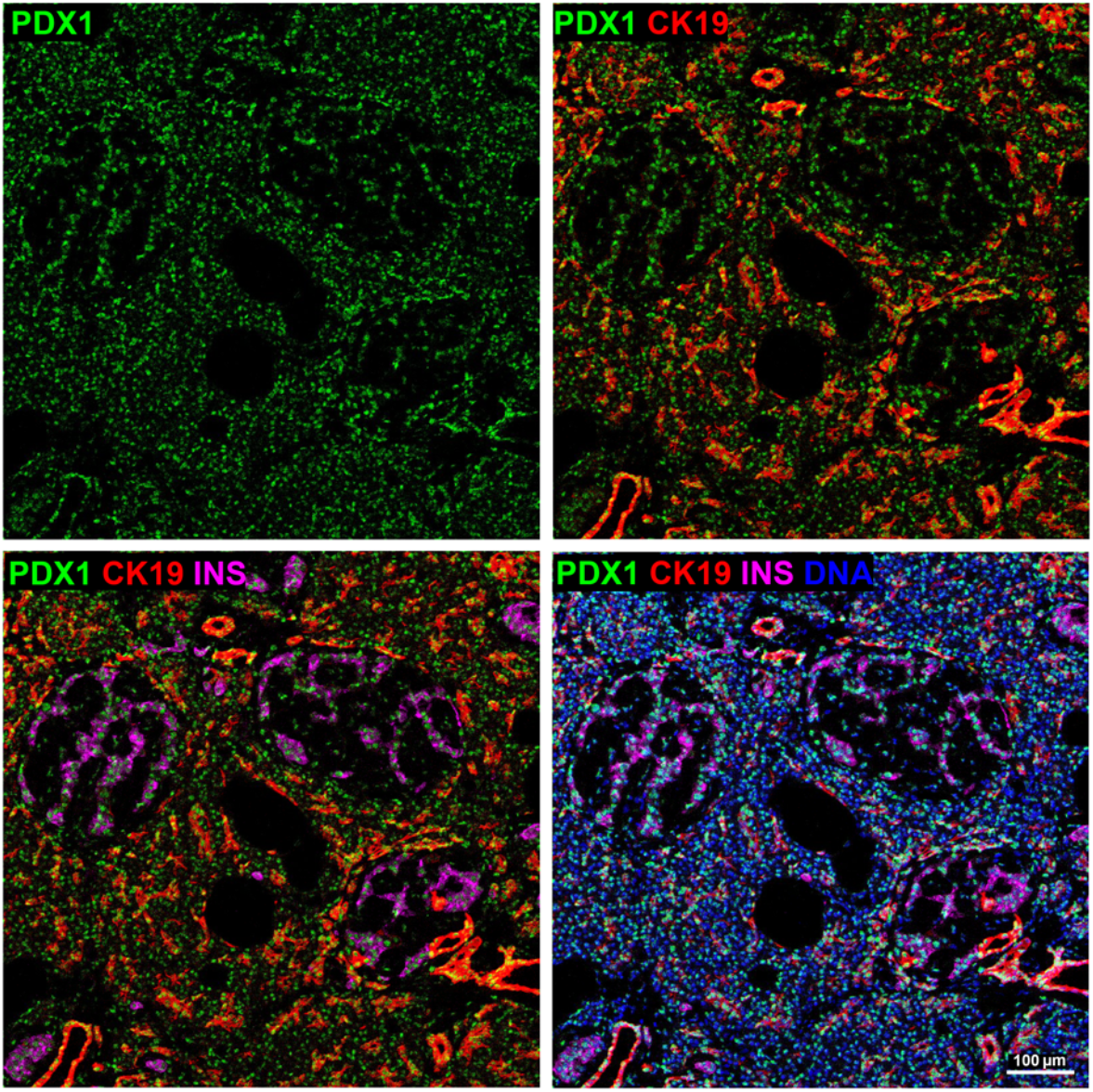
IMC staining of PDX1 in the pancreas tissue. PDX1 were expressed both in islet beta cells and in ductal cells. Red, CK19; Green, PDX1; Magenta, INS; Blue, DNA.

**Figure S5.**
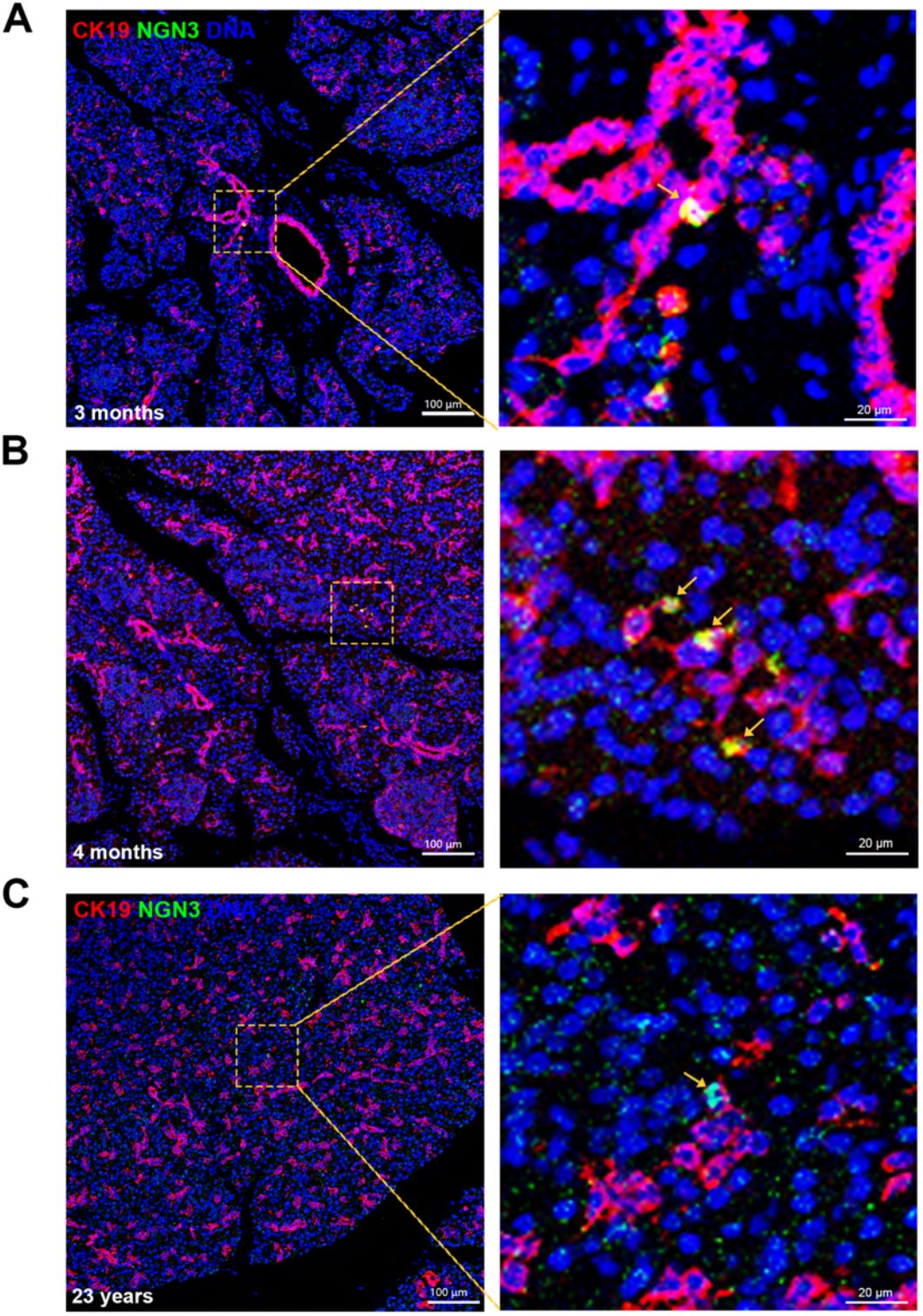
NGN3-positive ductal cells were observed in the infancy pancreases (A&B) and a young adult pancreas (C). Red, CK19; Green NGN3; Blue, DNA.

**Table S1.** Donor information.

| Donor No. | Age (years) | Sex | BMI (kg/m<sup>2</sup>) | HbA1c (%) |
| --- | --- | --- | --- | --- |
| HP060 | 0.08 | Female | 16.00 | / |
| HP067 | 0.25 | Female | 34.40 | / |
| HP220 | 0.29 | male | 16.53 | / |
| HP117 | 0.33 | Female | 24.69 | / |
| HP216 | 0.58 | male | 16.33 | / |
| HP038-ch | 1 | Female | 8.33 | / |
| HP217 | 2 | male | 13.58 | / |
| HP114 | 4.08 | Female | 15.00 | / |
| HP225 | 7 | Female | 15.63 | / |
| HP057 | 8 | male | 14.79 | / |
| HP190 | 8 | Female | 16.38 | / |
| HP222 | 8 | male | 14.29 | / |
| HP211 | 9 | Female | 15.31 | / |
| HP092 | 10 | male | 16.65 | 5.6 |
| HP137 | 12 | male | / | / |
| H150 | 13 | male | / | / |
| HP201 | 13 | male | / | / |
| HP041 | 14 | male | 19.03 | 4.9 |
| HP070 | 14 | male | 20.20 | 5.7 |
| HP082 | 14 | male | 21.48 | 5.2 |
| HP083 | 16 | male | 20.20 | 4.1 |
| HP130 | 17 | male | 20.28 | 5 |
| HP069 | 18 | male | 19.59 | 4.2 |
| HP089 | 20 | male | 20.05 | 5.6 |
| H109 | 23 | male | 22.49 | 4.7 |
| H123 | 25 | male | 19.59 | 5.2 |
| HP129 | 25 | Female | 21.48 | 5 |
| HP165 | 27 | male | 22.86 | 5.5 |
| HP155 | 29 | male | 23.03 | 4.3 |
| HP051 | 30 | male | 23.66 | 5.5 |
| HP025 | 37 | male | 21.72 | 5.5 |
| HP028 | 44 | male | 22.49 | 4.3 |
| HP035 | 45 | male | 27.78 | 5.6 |
| HP111 | 47 | male | 27.34 | 5.4 |
| HP039 | 49 | Female | 30.82 | 5.5 |
| HP046 | 52 | male | 20.76 | 5.1 |
| HP052 | 53 | Female | 20.96 | 5.5 |
| HP048 | 57 | male | 22.49 | 5.5 |
| HP045 | 58 | male | 26.12 | 5.3 |
| HP034 | 43 | Female | 21.48 | 6.1 |
| HP112 | 44 | male | 23.66 | 5.9 |
| HP113 | 44 | male | 29.39 | 6.1 |
| HP061 | 53 | male | 23.51 | 6 |
| HP050 | 60 | male | 22.86 | 5.9 |
| HP037 | 42 | male | 30.86 | 6.3 |
| HP019 | 44 | male | 24.20 | 6.1 |
| HP053 | 45 | male | 26.23 | 6.5 |
| HP018 | 47 | male | 32.40 | 6.7 |
| HP014 | 49 | male | 29.40 | 6.9 |
| HP017 | 49 | male | 24.20 | 7.6 |
| HP099 | 51 | male | 22.49 | 11.9 |
| HP038 | 55 | male | 22.59 | 6.8 |
| HP105 | 57 | Female | 24.97 | 5.8 |
| HP027 | 59 | male | 22.49 | 9.6 |
| HP012 | 63 | male | 25.95 | 7.2 |
| HP055 | 64 | Female | 23.43 | 8.9 |
| HP054 | 65 | male | 23.93 | 7.5 |
| HP006 | 60 | male | 30.93 | 6.4 |
| HP145 | 61 | male | 27.89 | 5.7 |
| HP149 | 61 | male | 32.87 | 5.2 |
| HP169 | 63 | male | 24.22 | 5 |
| HP110 | 65 | Female | 21.48 | 5.8 |
| HP148 | 66 | male | 25.71 | 5.8 |
| HP062 | 67 | male | 25.06 | 4.4 |
| HP178 | 68 | male | 22.86 | / |
| HP119 | 69 | Female | 18.73 | / |
| HP056 | 73 | Female | 25.71 | 5.4 |

**Table S2.**
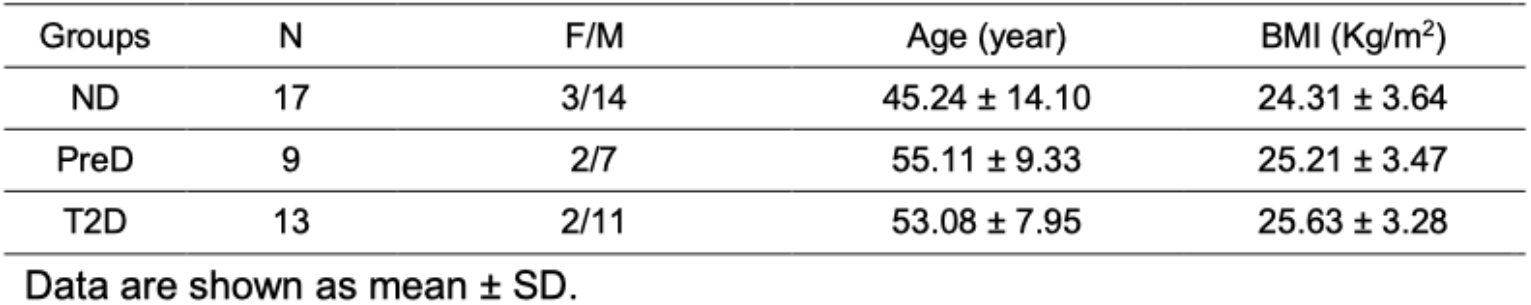
Donor information in ND, PrnD, T2D groups.

**Table S3.** Antibody information.

| Antibodies | Source | Cat. |
| --- | --- | --- |
| Insulin | Abcam | ab63820 |
| NKX6.1 | Novus | NBP1-49672 |
| PDX1 | CST | D59H3 |
| UCN3 | Sigma | HPA038281 |
| Glucagon | Abcam | ab10988 |
| ARX | R&D | AF7068 |
| Somatostatin | Santa Cruz | sc-55565 |
| Pancreatic Polypeptide | Abcam | ab77192 |
| Synaptophysin | Abcam | ab32127 |
| ALDH1a3 | Abcam | NBP2-15339 |
| NGN3 | R&D | AF3444 |
| SOX9 | Abnova | H00006662-M02 |
| CK19 (cytokeratin 19) | Novus | NB100-687SS |
| CD34 | Abcam | ab81289 |
| α-SMA | Sigma | A5228 |
| Ki67 | Abcam | ab15580 |
| Histone H3 Phospho (Ser28) | BioLegend | 641002 |
| GHRELIN | Santa Cruz | sc-293422 |
| CD99 | R&D | AF3968 |
| β-actin | CST | 3700s |

